# Conformational Change of Syntaxin-3b in Regulating SNARE Complex Assembly in the Ribbon Synapses

**DOI:** 10.1101/2021.09.07.459295

**Authors:** Claire Gething, Joshua Ferrar, Bishal Misra, Giovanni Howells, Ucheor B. Choi

**Affiliations:** Department of Biochemistry, West Virginia University

**Keywords:** syntaxin-3b, SNARE complex assembly, Munc18-1, single-molecule FRET

## Abstract

Neurotransmitter release of synaptic vesicles relies on the assembly of the soluble N-ethylmaleimide-sensitive factor attachment protein receptor (SNARE) complex, consisting of syntaxin and SNAP-25 on the plasma membrane and synaptobrevin on the synaptic vesicle. The formation of the SNARE complex progressively zippers towards the membranes, which drives membrane fusion between the plasma membrane and the synaptic vesicle. However, the underlying molecular mechanism of SNARE complex regulation is unclear. In this study, we investigate the syntaxin-3b isoform found in the retinal ribbon synapses using single-molecule fluorescence resonance energy transfer (smFRET) to monitor the conformational changes of syntaxin-3b that modulate the SNARE complex formation. We found that syntaxin-3b is predominantly in a self-inhibiting closed conformation, inefficiently forming the ternary SNARE complex. Conversely, a phosphomimetic mutation (T14E) at the N-terminal region of syntaxin-3b promoted the open conformation, similar to the constitutively open form of syntaxin LE mutant. When syntaxin-3b is bound to Munc18-1, SNARE complex formation is almost completely blocked. Surprisingly, the T14E mutation of syntaxin-3b partially abolishes Munc18-1 regulation, acting as a conformational switch to trigger SNARE complex assembly. Thus, we suggest a model where the conformational change of syntaxin-3b induced by phosphorylation initiates the release of neurotransmitters in the ribbon synapses.

## Introduction

Neurotransmission in the retina is an essential process of vision, where light captured by photoreceptors is transduced into electrical signals to modulate the release of neurotransmitters at specialized ribbon synapses. Ribbon synapses contain a synaptic ribbon, which is a large, proteinaceous, horseshoe-shaped structure anchored to the plasma membrane and tethered by many synaptic vesicles on the surface ^1–3^. Similar to conventional synapses, neurotransmitters are released at the active zone through a Ca^2+^ dependent exocytosis of synaptic vesicles within the plasma membrane ^4^. However, ribbon synapses release neurotransmitters in a slow and graded manner ^1–3^. The presynaptic active zone contains essential fusion machineries such as the soluble N-ethylmaleimide-sensitive factor attachment receptor (SNARE) proteins ^5–7^. Conventional SNAREs consist of SNAP-25 and syntaxin-1 on the plasma membrane and synaptobrevin-2 on the synaptic vesicle, forming a stable four-helix bundle, i.e., the SNARE complex, which allows the synaptic vesicle and the plasma membrane to be in close proximity, permitting membrane fusion ^8, 9^. The ribbon synapses that reside in the retina also contain SNAP-25 and synaptobrevin-2, yet lack syntaxin-1. Instead, it has been shown that the ribbon synapses are mainly localized with the syntaxin-3b isoform ^10–13^.

Genetic studies have identified four different isoforms of syntaxin-3 (syntaxin-3a, 3b, 3c, and 3d) which have been generated by splice variants of the mouse syntaxin-3 gene ^13^. Syntaxin-3a and -3b have identical N-terminus consisting of an N-terminal peptide, a three α-helical bundle called the Habc domain, and a linker region, but differ in the SNARE motif and the C-terminal transmembrane domain. Syntaxin-3c and -3d do not contain the SNARE motif and the transmembrane domain, and therefore are unlikely to form the ternary SNARE complex. Structural studies have identified syntaxin-1a in two conformations. The first is where the Habc domain of syntaxin-1a folds back to the SNARE motif in a closed conformation bound to sec1/Munc18-like (SM) protein Munc18-1, and the second is the open conformation of syntaxin-1a, forming ternary SNARE complex with SNAP-25 and synaptobrevin-2 ^14–18^. Despite the similar domain structure between syntaxin-1a and syntaxin-3b, the latter was observed to be less fusogenic, illustrated using a reconstituted membrane fusion assay ^13^. This property is due to the lower binding affinity of SNAP-25 to syntaxin-3b when compared to syntaxin-1a. Later, studies found that phosphorylation at residue 14 of syntaxin-3b by the Ca^2+^/calmodulin-dependent protein kinase II (CaMKII) increased the binding with SNAP-25, promoting efficient assembly of binary t-SNARE complex ^12^.

Munc18-1 is a SNARE chaperone that helps facilitate SNARE complex formation as well as regulate the process via inhibiting SNARE complex assembly by locking syntaxin-1a in a closed conformation. The closed conformation of syntaxin-1a by Munc18-1 is later catalyzed by the priming factor Munc13-1 to transition to the open conformation, allowing SNARE complex formation ^15, 19–21^. A mutation in the linker region connecting the Habc and the SNARE motif, referred to as the LE mutant (L165A, E166A), was found to bypass the requirement of Munc13-1 to transition from the closed conformation, locked by Munc18-1, to the open conformation, permitting ternary SNARE complex formation when SNAP-25 and synaptobrevin are introduced *in vitro* ^15^.

Here, we applied single-molecule fluorescence resonance energy transfer (smFRET) to directly monitor the conformational dynamics of syntaxin-3b during SNARE complex formation. smFRET is capable of distinguishing heterogeneous populations and intermediates of protein conformations, which is typically lost in ensemble measurements due to averaging behavior. We found that almost 90 % of syntaxin-3b molecules are in the closed conformation, preventing binary t-SNARE complex formation with SNAP-25. The closed population decreased to about 50 % when a phosphomimetic mutation (T14E) was introduced and is further decreased in the LE mutation of syntaxin-3b. Even in the presence of Munc18-1, both the syntaxin-3b T14E mutant and LE mutant were unable to transition back to the closed conformation, permitting binary t-SNARE complex formation with SNAP-25. This agrees with previous findings where phosphorylation operates as a conformational switch of syntaxin-3b to promote SNARE complex assembly.

## Results

### Syntaxin-3b is predominantly in the closed conformation

Like many other syntaxins, syntaxin-3b consists of an N-terminal peptide, the Habc domain, a linker region, the SNARE motif, and a transmembrane region (Fig. 1A) ^22^. The structure of full-length syntaxin-1a has been difficult to determine due to the unstructured linker region between the N-terminal Habc domain and the SNARE motif, permitting ensemble native state conformations. However, syntaxin-1a adopts a closed conformation when bound to Munc18-1, where the Habc domain folds back to the SNARE motif ^14, 15, 23, 24^. In addition, both syntaxin-1a and syntaxin-3b have similar disorder tendencies of 68% and 67%, respectively, when assessed via sequence analysis (Fig. 1A) ^25^. Despite the similarity between syntaxin-1a and syntaxin-3b, previous biochemical and functional studies found that syntaxin-3b inefficiently forms a complex with the t-SNARE SNAP-25 and was observed to be less fusogenic in a reconstituted vesicle fusion assay when compared to vesicles consisting of syntaxin-1a ^13^.

**Figure 1.**
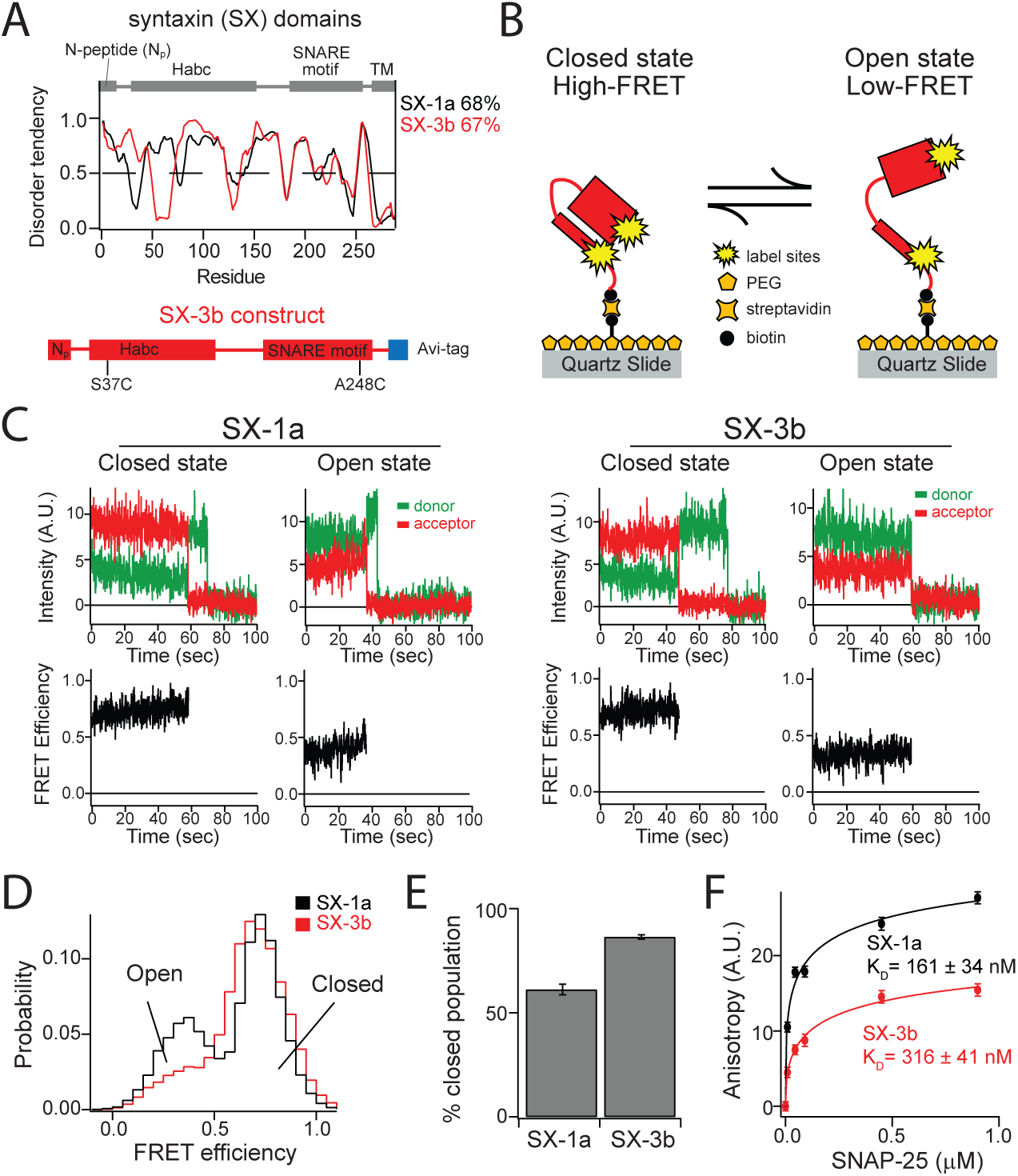
Native state conformations of syntaxin-1a and syntaxin-3b. **(A)** The disorder tendencies of syntaxin-1a and -3b are analyzed by PONDR VLXT ^25^. The protein domain diagram illustrates the common domains of syntaxin-1a and syntaxin-3b (top panel) and the syntaxin-3b construct used in this experiment (bottom panel). **(B)** Schematic of a surface-tethered syntaxin molecule labeled with FRET dye pairs on a functionalized surface of a microscope slide in a closed and open conformation leading to high and low FRET efficiency, respectively. **(C)** Representative single-molecule fluorescence intensity time traces of sytnaxin-1a (SX-1a) and syntaxin-3b (SX-3b) in the closed and open conformations (top panels). The donor and acceptor intensities were converted to FRET efficiency time traces (bottom panels). **(D)** smFRET efficiency histograms of isolated SX-1a and SX-3b. **(E)** Percent closed populations of SX-1a and SX-3b were extracted from (D) by fitting two Gaussian functions to the FRET efficiency histograms. Shown are means ± SD (n=3). **(F)** Bulk fluorescence anisotropy measurements of interactions between Alexa 488 labeled syntaxins and unlabeled SNAP-25 at 0 μM, 0.009 μM, 0.045 μM, 0.090 μM, 0.450 μM, 0.900 μM concentrations. The anisotropy curves are fit with Hill equations to estimate the disassociation constant K_d_. Shown are means ± SD (n=3).

In order to understand the differences, we investigated the conformations of isolated syntaxin-3b using single-molecule FRET. Syntaxin-3b contains only two native cysteine residues in the transmembrane region, which was replaced with an Avi-tag sequence for *in vivo* biotinylation (Fig. 1A). We covalently attached donor and acceptor FRET label pairs on residues 37 and 247 of syntaxin-3b by introducing cysteine residues on a cysteine-free construct (Fig. 1B). For comparison, we labeled syntaxin-1a on residues 35 and 249, as conducted previously ^15, 21^. These label sites were engineered based on the crystal structure of the sytnaxin-1a/Munc18-1 heterodimer complex, so syntaxin-3b adopts high FRET efficiency in the closed conformation and low FRET efficiency in the open conformation. To achieve single-molecule resolution, we replaced the C-terminal transmembrane region with an Avi-tag sequence for surface tethering via a biotin-streptavidin linkage (Fig. 1B). The surface of the microscope slide was passivated with polyethylene glycol (PEG) to prevent non-specific binding of molecules on the surface and mimic the geometry of the presynaptic environment where syntaxins are localized on the plasma membrane via the C-terminal transmembrane region.

We measured the donor and acceptor fluorescent intensity time traces of individual molecules and calculated FRET efficiencies of sytnaxin-1a and syntaxin-3b in their native state conditions (Fig. 1C). Anti-correlation between donor and acceptor intensities indicates transitions between FRET efficiencies, i.e., the conformational states, or photobleaching of the acceptor dye molecule where the donor intensity rises with respect to the stepwise drop of the acceptor intensity (Fig. 1C top panels). All single molecule intensity time traces of syntaxin-1a and syntaxin-3b showed two distinct high and low FRET efficiencies prior to the acceptor or donor photobleaching event occurring. All single molecules, i.e., molecules with one donor and one acceptor dye molecule, were plotted using FRET efficiency histograms to determine the native state conformations. The distribution of FRET efficiencies for both syntaxin-1a and syntaxin-3b revealed two major populations at high and low FRET efficiencies indicating the closed and the open conformations, respectively (Fig. 1D). Two Gaussian functions were used to fit the FRET efficiency histograms to quantify the percent populations of each state.

Interestingly, almost 90% of syntaxin-3b was in the closed conformation compared to about 60% for syntaxin-1a (Fig. 1E). This is consistent with the previous expectation that syntaxin-3b would likely be in a more closed conformation compared to syntaxin-1a, ultimately lowering the affinity to bind SNAP-25 ^13^. To further corroborate the interaction of SNAP-25 with syntaxin-1a and syntaxin-3b, we conducted ensemble fluorescence anisotropy measurements to obtain an estimate of the dissociation constant (K_d_). Syntaxin-1a and syntaxin-3b were labeled at the primary amino group at the N-terminus with the fluorescent dye Alexa 488. The increase in anisotropy at varying concentrations of SNAP-25 indicates an increase in molecular mass, which is correlated with complex formation. The fluorescence anisotropy curves were fit to a Hill function, resulting in a K_d_ of about 161 nM and 316 nM for syntaxin-1a and syntaxin-3b, respectively (Fig. 1F). This is consistent with previous result of sytnaxin-1a and SNAP-25 binding affinity of 126 nM using a pulldown assay ^26^. This agrees well with our smFRET measurements where the increase in the closed conformation of syntaxin-3b results in an almost 2-fold decrease in the binding affinity compared to syntaxin-1a.

### Mutations altering the conformations of syntaxin-3b

Syntaxin-3b is enriched in the presynaptic plasma membrane of synaptic ribbon synapses and has been shown to be an essential component for synaptic vesicle exocytosis in the retinal bipolar cells ^10–13, 27, 28^. Despite the importance, *in vitro* binding assays demonstrated that syntaxin-3b reduces binding of SNAP-25, and a vesicle fusion assay showed reduced membrane fusion when reconstituted with syntaxin-3b compared to syntaxin-1a ^13^. Moreover, the removal of the Habc domain of syntaxin-3b increased the binding of SNAP-25, demonstrating that the differences in the SNARE motif of syntaxin-3b compared with syntaxin-1a were not the cause of reduced binding of SNAP-25 ^12^. Furthermore, smFRET measurements directly demonstrate that syntaxin-3b adopts a closed conformation in its native state, which is likely the cause of inhibiting SNARE complex formation with SNAP-25 and synaptobrevin-2 (Fig.1). Thus, the SNARE-mediated membrane fusion of synaptic vesicles requires other mechanisms to allow the opening of syntaxin-3b in order to promote the release of neurotransmitters efficiently.

Previously, phosphorylation on syntaxins has been shown to increase sustained synaptic vesicle exocytosis in hippocampal neuronal cultures ^29–31^. Sytnaxin-3b has also been found to be a substrate for Ca^2+^/calmodulin-dependent protein kinase II (CaMKII) at residue 14 ^12^. Therefore, we investigated the effect of phosphorylation on the conformations of syntaxin-3b using smFRET by introducing a T14E phosphomimetic mutation (Fig. 2A). For comparison, we also examined the well-known LE mutation (L165A, E166A) on sytnaxin-3b, which has been found to induce the open conformation of sytnaxin-1a and bypass the requirement of the priming factor Munc13-1 when bound with Munc18-1 in a self-inhibiting closed conformation (Fig. 2A) ^15–17^. Syntaxin-3b mutants were labeled with donor and acceptor FRET dye pairs and surface tethered via a biotin-streptavidin linkage at the C-terminus of syntaxin-3b on a passivated surface of the microscope slide (Fig. 2B). Interestingly, compared to wild-type (WT) syntaxin-3b, where stable high FRET efficiencies were observed, the syntaxin-3b T14E and the LE mutants show stochastic transitions between high and low FRET efficiencies (Fig. 2C). All single molecules, i.e., molecules with FRET pairs consisting of one donor and one acceptor dye molecule, were plotted in FRET efficiency histograms to determine the conformational states of sytnaxin-3b mutants. For both the T14E and LE mutants, the major population in the FRET efficiency histogram shifted to the low FRET efficiency (Fig. 2D). The histograms were quantified by fitting two Gaussian functions to determine the percent closed population. Interestingly, the T14E phosphomimetic mutation on syntaxin-3b decreased the percent closed population to almost 50% whereas the LE mutation nearly abolished the closed conformation of sytnaxin-3b (Fig. 2E). Since the opening of sytnaxin-3b should induce binary t-SNARE complex formation with SNAP-25 by increasing the binding affinity, we conducted ensemble fluorescence anisotropy measurements by labeling syntaxin-3b mutants with Alexa 488 at varying concentrations of SNAP-25. In line with the opening of sytnaxin-3b, the T14E and the LE mutant increased the affinity of SNAP-25 with an estimated K_d_ of 171 nM and 76 nM, respectively (Fig. 2F).

**Figure 2.**
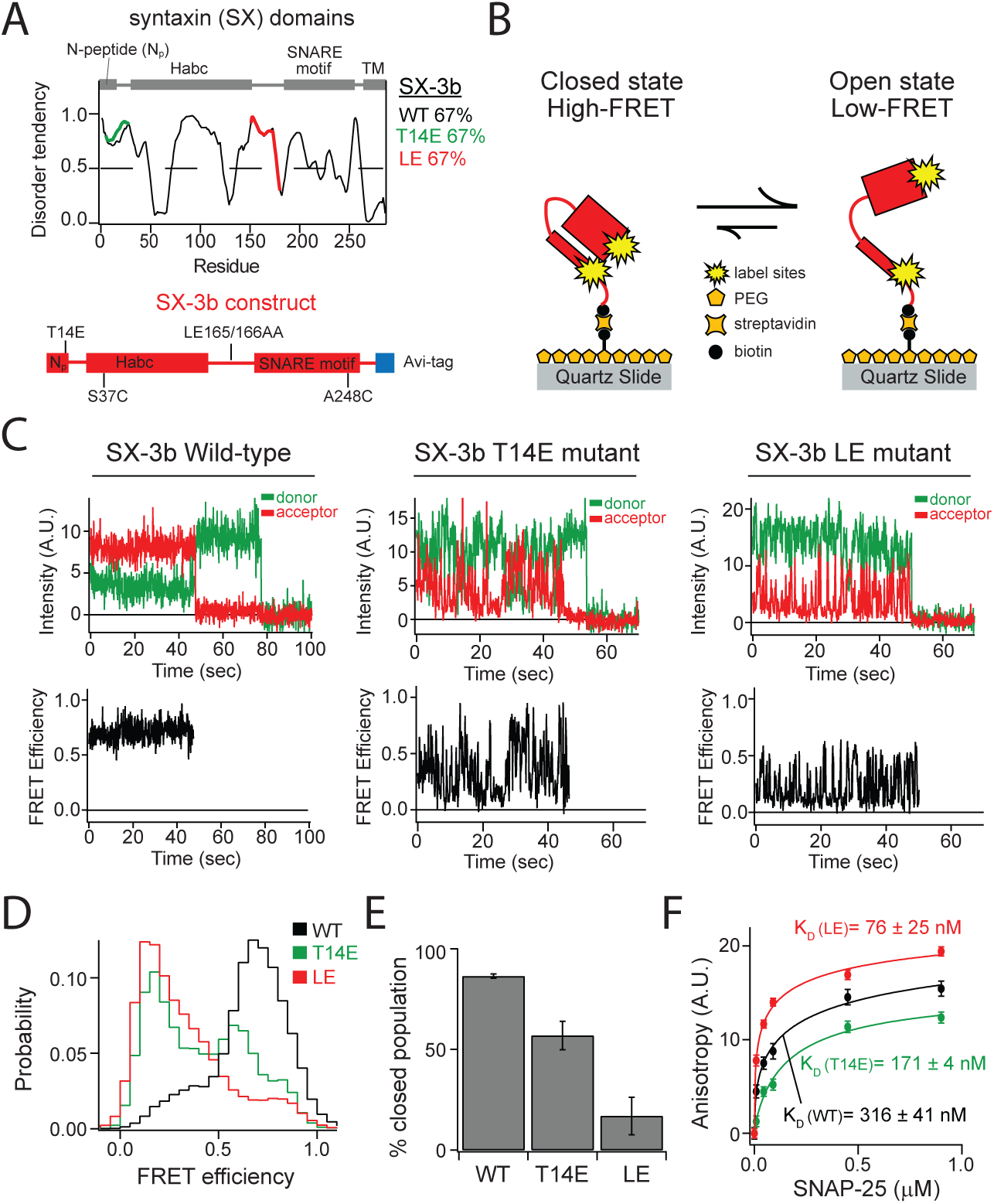
The phosphomimetic T14E and the LE mutations alter the conformation of syntaxin-3b. **(A)** The disorder tendencies of WT, T14E, and LE (L165A, E166A) mutation of syntaxin-3b are analyzed by PONDR VLXT ^25^. The protein domain diagram illustrates the common domains in syntaxin-1a and syntaxin-3b (top panel) and the syntaxin-3b construct used in this experiment (bottom panel). **(B)** Schematic of a surface-tethered syntaxin-3b molecule labeled with FRET dye pairs on a functionalized surface of the microscope slide in a closed and open conformation leading to high and low FRET efficiency, respectively. **(C)** Representative single-molecule fluorescence intensity time traces of sytnaxin-3b wild-type (WT) (left panels), T14E (middle panels), and LE (right panels) mutants. The donor and acceptor intensities were converted to FRET efficiency time traces (bottom panels). **(D)** smFRET efficiency histograms of WT, T14E and LE syntaxin-3b mutants. **(E)** Percent closed populations of syntaxin-3b WT, T14E, and LE mutant were extracted from (D) by fitting two Gaussian functions to the FRET efficiency histograms. Shown are means ± SD (n=3). **(F)** Bulk fluorescence anisotropy measurements of interactions between Alexa 488 labeled syntaxins and unlabeled SNAP-25 at 0 μM, 0.009 μM, 0.045 μM, 0.090 μM, 0.450 μM, 0.900 μM concentrations. The anisotropy curves are fit with Hill equations to estimate the disassociation constant K_d_. Shown are means ± SD (n=3).

### Phosphorylation of sytnaxin-3b promotes SNARE complex formation

During SNARE complex formation, the SNARE motifs of syntaxin-1a, SNAP-25, and synaptobrevin-2 assemble into a four-helical bundle, where two helices are derived from SNAP-25 ^8, 32^. Here, we conducted a smFRET assay to monitor the conformational changes of syntaxin-3b and its mutants during SNARE complex assembly with SNAP-25 and synaptobrevin-2 (Fig.3A). Wild-type syntaxin-3b and its mutants were labeled with donor and acceptor FRET dye pairs, and smFRET measurements were conducted in the presence of SNAP-25 and the cytoplasmic domain of synaptobrevin-2 (residues 1–96) (Fig. 3B). In addition, FRET labeled syntaxin-3b molecules were surface tethered on a passivated surface of the microscope slide via a biotin-streptavidin linkage, and 10 μM of SNAP-25 and synaptobrevin-2 were injected prior to data collection.

**Figure 3.**
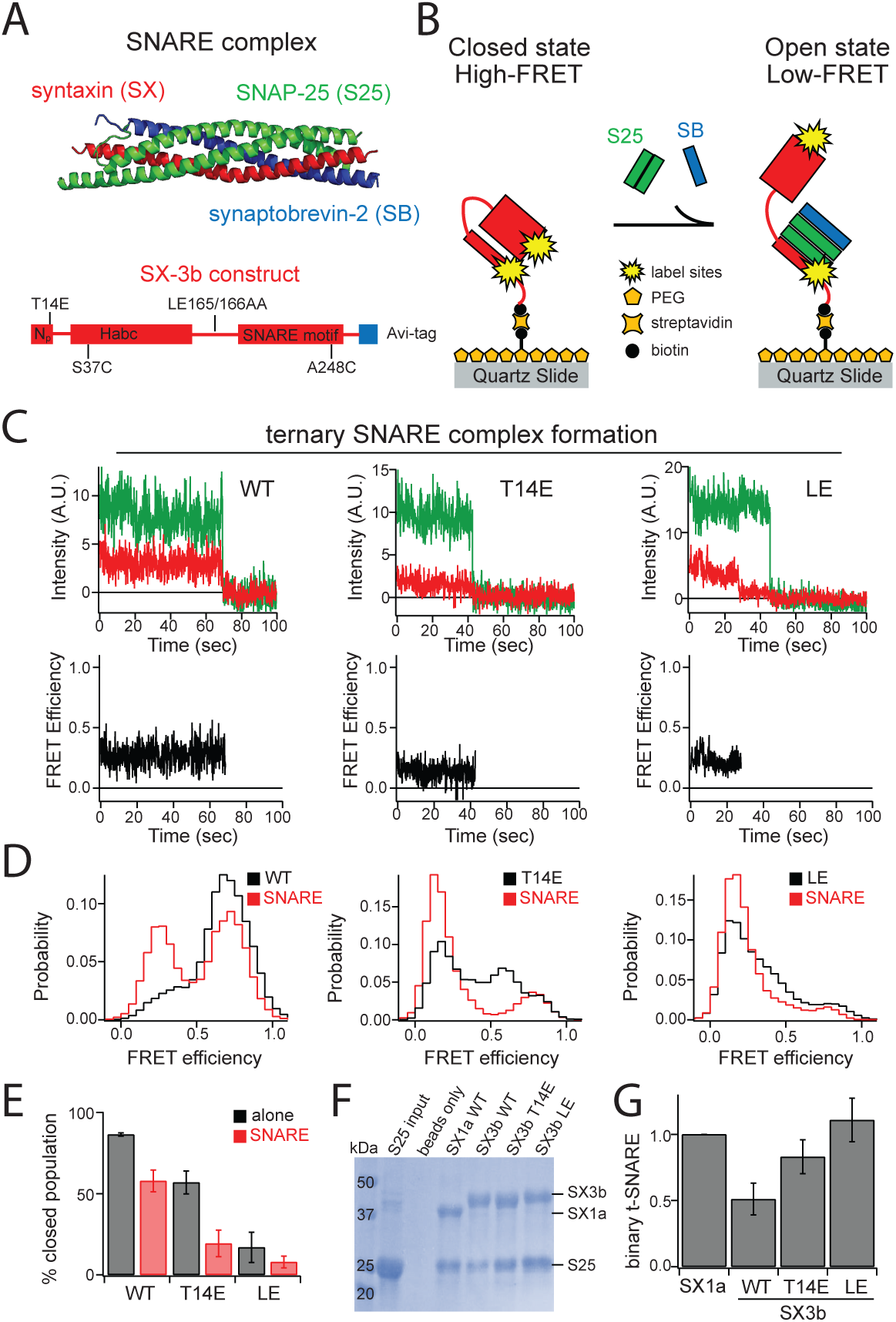
SNARE complex formation of syntaxin-3b and its mutants. **(A)** The structure of the SNARE complex forming a four-helical bundle (PDB ID: 1SFC). Protein domain diagram of syntaxin-3b constructs used in this experiment (bottom panel). **(B)** Schematic of a surface-tethered syntaxin-3b molecule labeled with FRET dye pairs on a functionalized surface of the microscope slide in a closed conformation and the transit to an open conformation during SNARE complex assembly in the presence of 10 μM SNAP-25 and synaptobrevin-2 in solution. **(C)** Representative single-molecule fluorescence intensity time traces of wild-type (left panels), T14E (middle panels) and LE (right panels) mutations on syntaxin-3b in the presence of 10 μM SNAP-25 and synaptobrevin-2 in solution. The donor and acceptor intensities were converted to FRET efficiency time traces (bottom panels). **(D)** smFRET efficiency histograms of wild-type, T14E and LE mutation on syntaxin-3b in the presence of 10 μM SNAP-25 and synaptobrevin-2 in solution. **(E)** Percent closed populations of syntaxin-3b WT, T14E, and LE mutant in the presence of 10 μM SNAP-25 and synaptobrevin-2 in solution were extracted from (D) by fitting two Gaussian functions to the FRET efficiency histograms. Shown are means ± SD (n=3). **(F–G)** A pulldown assay was conducted using biotinylated syntaxins as bait on neutravidin coated beads to estimate the efficiency of binary t-SNARE complex formation with SNAP-25. Proteins bound to the beads were confirmed by SDS-PAGE. SNAP-25 bands, indicating the binary t-SNARE complex formation, were analyzed using ImageJ software (NIH, Bethesda, MD). Shown are means ± SD (n=3).

As noted in the intensity time traces of syntaxin-3b and its mutants showing stable FRET efficiencies prior to photobleaching events, the major peak of the smFRET distributions shifted to a low FRET efficiency (Fig. 3C–D). This indicates that syntaxin-3b formed SNARE complex, resulting in the opening of the Habc domain of syntaxin-3b. The smFRET distributions were fit with two Gaussian functions to quantify the closed populations. In the presence of SNAP-25 and synaptobrevin-2, the percent of closed populations for all syntaxin-3b mutants decreased, which correlates to about a 3-fold increase in the open populations for syntaxin-3b WT and T14E mutant (Fig. 3E). The closed population for syntaxin-3b LE mutant was similar in the presence of SNAP-25 and synaptobrevin-2 since the LE mutation initially induced an open conformation of syntaxin-3b.

Following the smFRET experiments, we further verified the conformational opening of syntaxin-3b mutants by conducting a pulldown assay to monitor the binary t-SNARE complex formation with SNAP-25 (Fig. 3F). Purified syntaxin proteins were used as a bait via biotinylation at the N-terminal primary amino group and subsequently binding to neutravidin-coated beads. As a negative control, neutravidin coated beads alone were used to confirm that proteins do not non-specifically bind to the beads. SNAP-25 was incubated for 1 hour at room temperature following a rinsing step with buffer before the bead samples were collected for gel analysis. The intensities of the protein bands for SNAP-25 were normalized based on the correlating syntaxin bands. Compared to syntaxin-1a, approximately 50% less SNAP-25 formed binary t-SNARE complex with WT syntaxin-3b (Fig. 3F–G). The amount of binary t-SNARE complex formation increased about 30% with the syntaxin-3b T14E mutation. On the other hand, the syntaxin-3b LE mutant dramatically increased the binding of SNAP-25, indicating that the LE mutation induces the open conformation of syntaxin-3b to efficiently form the binary t-SNARE complex.

### Munc18-1 reverts syntaxin-3b phosphomimetic mutant back to the closed conformation inhibiting SNARE complex formation

In addition to SNAREs, membrane fusion of synaptic vesicles requires Sec1/Munc18-family (SM) protein Munc18-1 ^5^. Knockout of Munc18-1 completely abolishes neurotransmitter release in mice ^33^. Moreover, Munc18-1, along with the priming factor Munc13-1, orchestrates SNARE complex assembly by preventing disassembly via the ATPase, NSF ^20, 34^. First, Munc18-1 locks syntaxin-1a in a self-inhibiting closed conformation, acting as an acceptor complex for synaptobrevin-2 and the priming factor Munc13-1 to interact with ^35^. Moreover, recent studies discovered a new function of Munc18-1, where it cooperates with Munc13-1 to ensure proper assembly of the SNARE complex, promoting efficient membrane fusion in reconstituted systems ^21^. Munc18-1 is therefore a key regulator that is capable of both inhibiting and catalyzing SNARE complex assembly.

Removal of the N-peptide of syntaxin-1a, i.e., deleting the N-terminal phosphorylation site, was capable of facilitating SNARE complex assembly even in the presence of Munc18-1 ^23^. Additionally, the LE mutation on syntaxin-1a successfully formed SNARE complex in the presence of Munc18-1 despite the strong binding affinity and bypassing the requirement of the priming factor Mun13-1 ^15, 23^. However, recent X-ray crystal structure and small-angle X-ray scattering (SAXS) data studies of syntaxin-1a lacking the N-peptide or with the LE mutation was observed in a closed conformation when bound to Munc18-1 ^24^. Therefore, to understand the effect of the SNARE chaperone Munc18-1 on syntaxin-3b, we conducted smFRET measurements to directly monitor the consequences of Munc18-1 on syntaxin-3b and its mutants (Fig. 4A–B). Similar as before, we surface tethered donor and acceptor FRET pair labeled sytnaxin-3b and its mutants on a passivated surface of the microscope slide via a biotin-streptavidin linkage and conducted smFRET measurements in the presence of 1 μM Munc18-1 in solution (Fig. 4B). The intensity time traces of syntaxin-3b and its mutants in the presence of Munc18-1 were converted to FRET efficiencies and plotted in histograms to determine the conformational states induced by Munc18-1 (Fig. 4C–D). Since syntaxin-3b is predominantly in the closed conformation, there was little impact from Munc18-1 as observed by the FRET efficiency histogram of wild-type syntaxin-3b in the presence of Munc18-1 (Fig. 4D–E). However, the major population of the FRET efficiency histogram of the phosphomimetic T14E mutant of syntaxin-3b shifted to high FRET efficiency in the presence of Munc18-1 (Fig. 4D middle panel, and Fig. 4E). This is in agreement with previous structural studies where syntaxin-1a lacking the N-peptide was observed in the closed conformation bound to Munc18-1 ^24^. Interestingly, the syntaxin-3b LE mutant in the presence of Munc18-1 slightly increased to mid FRET efficiency, but was unable to recover to the high FRET efficiency, i.e., the closed conformation (Fig. 4C right panel, and Fig. 4E).

**Figure 4.**
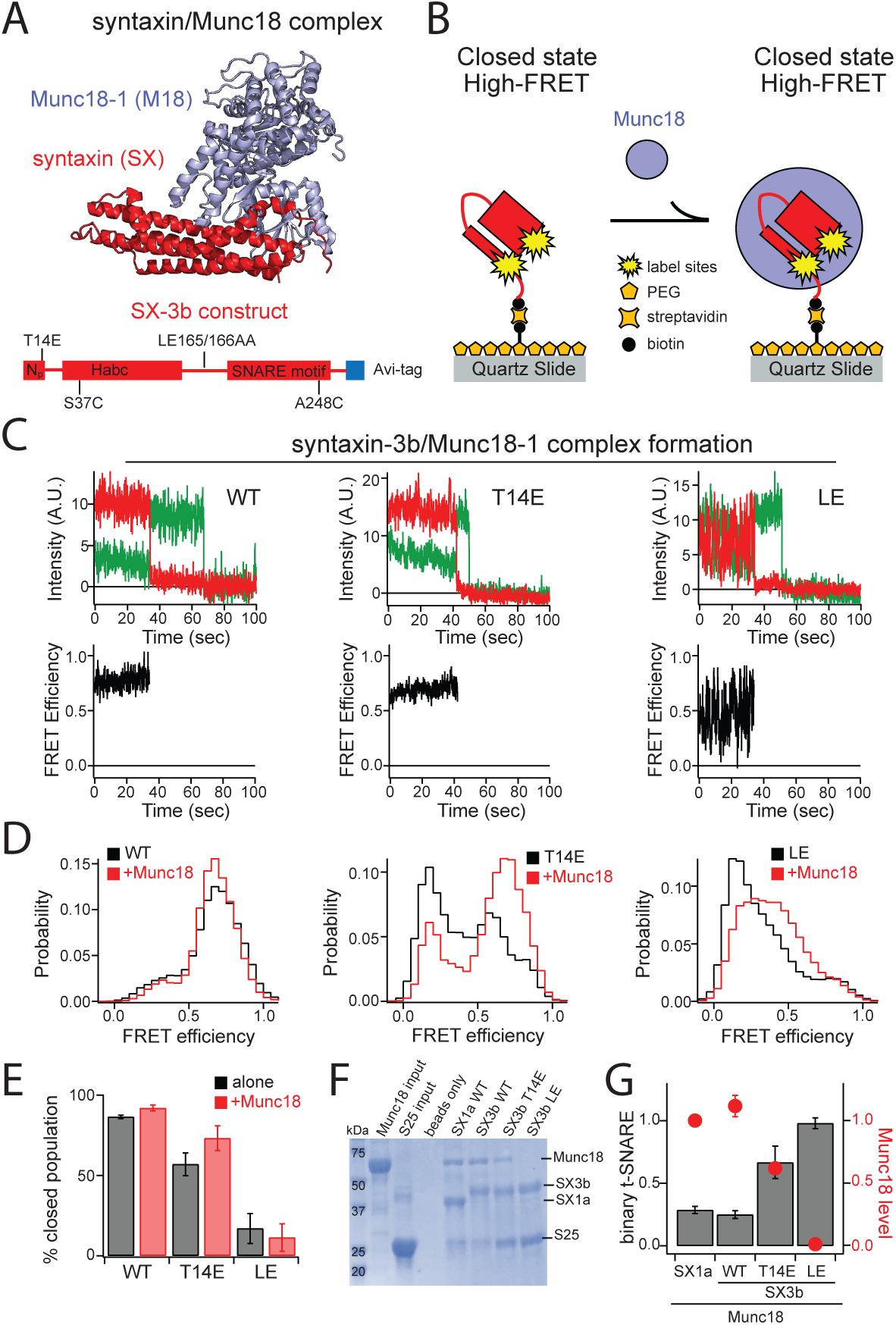
Effect of Munc18-1 on the conformation of syntaxin-3b and its mutants. **(A)** The structure of syntaxin/Munc18 complex showing the closed conformation of syntaxin (PDB ID: 3C98). Protein domain diagram of syntaxin-3b constructs used in this experiment (bottom panel). **(B)** Schematic of a surface-tethered syntaxin-3b molecule labeled with FRET dye pairs on a functionalized surface of the microscope slide in a closed conformation and locked in a closed conformation when bound to 1 μM Munc18-1 in solution. **(C)** Representative single-molecule fluorescence intensity time traces of WT (left panels), T14E (middle panels) and LE (right panels) mutation on syntaxin-3b in the presence of 1 μM Munc18-1 in solution. The donor and acceptor intensities were converted to FRET efficiency time traces (bottom panels). **(D)** smFRET efficiency histograms of wild-type, T14E and LE mutations on syntaxin-3b in the presence of 1 μM Munc18-1 in solution. **(E)** Percent closed populations of syntaxin-3b WT, T14E, and LE mutant in the presence of 1 μM Munc18-1 in solution were extracted from (D) by fitting two Gaussian functions to the FRET efficiency histograms. Shown are means ± SD (n=3). **(F–G)** A pulldown assay was conducted using biotinylated syntaxins in complex with Munc18-1 as bait on neutravidin coated beads to estimate the efficiency of binary t-SNARE complex formation with SNAP-25. Proteins bound to the beads were confirmed by SDS-PAGE. SNAP-25 bands, indicating the binary t-SNARE complex formation, as well as Munc18-1 bands were analyzed using ImageJ software (NIH, Bethesda, MD). Shown are means ± SD (n=3).

In addition to the smFRET measurements, we conducted pulldown assays using syntaxin/Munc18-1 complex as bait to test the efficiency of binary t-SNARE complex formation with SNAP-25 (Fig. 4F). Similar to Fig. 3F, purified syntaxins were bound to neutravidin-coated beads via biotin labeling at the primary N-terminal amine group of each syntaxin. For negative control, neutravidin-coated beads alone were used to confirm that proteins do not non-specifically bind to the beads. Different from previous pulldown assay, Munc18-1 was incubated for 1 hour at room temperature following a rinsing step with buffer before SNAP-25 was added to form the binary t-SNARE complex. After 1 hour incubation of SNAP-25 to syntaxin/Munc18 coated beads, the bead samples were collected for gel analysis after rinsing the beads with TBS buffer. The protein bands were analyzed using ImageJ software and the intensities for Munc18-1 and SNAP-25 bands were normalized based on the intensity of the correlating syntaxin bands. Munc18-1 bound tightly to syntaxin-1a and WT syntaxin-3b, however, the Munc18-1 level decreased to about 62% with the T14E mutation on syntaxin-3b (Fig. 4F–G). This is consistent with the notion that the T14E on syntaxin-3b induces an open conformation, therefore allowing efficient binary t-SNARE complex formation in the presence of Munc18-1 assessed by the pulldown assay (Fig. 4F–G). Moreover, Munc18-1 completely dissociated from syntaxin-3b LE mutant, promoting complete recovery of the binary t-SNARE complex formation (Fig. 4F–G). This agrees with the smFRET measurements of the syntaxin-3b LE mutant, where there was no increase in the closed population, i.e., the high-FRET state, in the presence of 1 μM Munc18-1 in solution (Fig. 4D right panel).

## Discussion

Exocytosis at the presynaptic terminal relies on a set of SNARE proteins to drive membrane fusion between the synaptic vesicle and the plasma membrane ^5, 22, 36, 37^. The core fusion machinery consists of the t-SNARE syntaxin and SNAP-25 on the plasma membrane and the v-SNARE synaptobrevin on the synaptic vesicle ^8^. Unlike the other SNAREs, the plasma membrane t-SNARE syntaxins (i.e., syntaxin 1–4) are composed of the regulatory N-terminal Habc domain, which causes sytnaxins to adopt two different conformations, i.e., an open and a closed conformation ^16, 38^. Prior to SNARE complex assembly, the plasma membrane syntaxins adopt a closed conformation tightly bound with the SNARE chaperone Munc18-1, where the syntaxin Habc domain folds back to the C-terminal SNARE motif ^5, 14^. However, recent structural studies showed that the yeast SNARE Tlg2 has been observed in an open conformation while in complex with the Munc18 homolog, Vps45 ^39^. The clamping mechanism of Munc18, locking syntaxin in a closed conformation, may be a specialized property in the neuronal SNAREs, which requires tight regulation and precise release of neurotransmitters. Therefore, during SNARE complex assembly, the syntaxin/Munc18 complex requires a brain-specific priming factor, such as Munc13-1, to catalyze the transition of the Habc domain of syntaxin to an open conformation, exposing the C-terminal SNARE motif and allowing for the initiation of SNARE complex formation in the presence of SNAP-25 and synaptobrevin ^15, 16, 18, 20^. Recent single-molecule fluorescence studies have demonstrated the conformation of isolated syntaxin-1a in a dynamic transition between an open and closed conformation, despite the complete closed conformation seen by NMR and SAXS data ^15, 17, 21, 24, 40, 41^. Despite the importance of the conformational switch of syntaxin in controlling synaptic vesicle exocytosis, the underlying molecular mechanism of other isoforms has not been investigated.

Here, we investigated the conformations of syntaxin-3b, which is a specialized SNARE protein for the exocytosis of synaptic vesicles in the ribbon synapses of the retina ^10, 13^. Different from syntaxin-1a in the conventional synapses of the nervous system, syntaxin-3b binds weakly to the t-SNARE SNAP-25, and an *in vitro* liposome fusion assay showed a decrease in the kinetics of membrane fusion when reconstituted with syntaxin-3b compared to syntaxin-1a ^13^. In spite of the overall sequence alignment showing 67.2% identity between syntaxin-1a and syntaxin-3b, the N-terminal half consisting of the Habc domain was less conserved at 56.8% identity than the SNARE motif at 75.9% identity (Supplemental Figure 1). Therefore, it is likely that the functional differences arise from the regulatory Habc domain of syntaxin-3b. It has been previously hypothesized that syntaxin-3b adopts a self-inhibiting closed conformation even without the SNARE chaperone Munc18-1, thus preventing SNARE complex assembly ^13^.

To directly monitor the conformations, we conducted single-molecule fluorescence measurements of isolated syntaxin-3b using smFRET. We found that isolated syntaxin-1a, located in the conventional synapses, adopts open and closed conformations with about 60% of the molecules in the closed population (Fig. 1). This is consistent with previous smFRET measurements where two conformations, i.e., an open and closed conformation, were observed for syntaxin-1a ^15, 21, 41^. Interestingly, almost 90% of isolated syntaxin-3b molecules were in the closed conformation (Fig. 1). Moreover, the affinity to SNAP-25 decreased by 2-fold when assessed by ensemble fluorescence anisotropy binding assay (Fig. 1F). Since the SNARE motifs of syntaxin-1a and syntaxin-3b are highly conserved, the decrease in the binding of SNAP-25 to syntaxin-3b is likely due to the interaction with the Habc domain folding back in a self-inhibiting closed conformation.

In conventional synapses in the brain, Munc18-1 acts as a molecular gatekeeper to lock syntaxin-1a in a closed conformation, preventing SNARE-mediated membrane fusion of synaptic vesicles and the plasma membrane ^5, 14^. Regardless of the tight heterodimeric complex of Munc18-1 and syntaxin-1a, the MUN domain of Munc13-1, a brain specific synaptic vesicle priming factor, is capable of catalyzing the transition of the closed conformation of syntaxin-1a latched with Munc18-1 to the open conformation for ternary SNARE complex formation ^15, 19, 20^. Deletion or mutation of Munc13s severely abolishes neurotransmitter release ^42–44^. In addition, Munc13-1/2 double-knockout mice showed complete loss of release-competent synaptic vesicles and severely altered the docking of synaptic vesicles ^45, 46^. In the retina, Munc13-2 is localized to the ribbon synapses ^47, 48^. Strikingly, deletion of Munc13-2 did not affect the synaptic signaling at the photoreceptor ribbon synapses and the readily releasable pool of docked synaptic vesicles, indicating that the ribbon synapse operates in a Munc13-independent manner ^47^. Moreover, the SNARE chaperone Munc18-1 is localized at the active zone of the ribbon synapses in the retina, regulating the opening of syntaxin-3b ^3^. This raises a question of how syntaxin-3b is released from the tight heterodimeric complex with Munc18-1 to promote SNARE complex assembly. Recent studies showed that syntaxin-3b is a substrate for Ca^2+^/calmodulin-dependent protein kinase II (CaMKII) and phosphorylates residue 14 at the N-terminus ^12^. Remarkably, the phosphomimetic mutation (T14E) of syntaxin-3b promoted binding to SNAP-25 when compared to the wild-type, indicating a novel mechanism of catalyzing the opening of syntaxin-3b in a calcium dependent phosphorylation by CaMKII ^12^. More recently, the phosphorylation at residue 14 of syntaxin-3b was demonstrated to be light regulated where the levels of phosphorylated syntaxin-3b significantly increased in the dark for rod photoreceptors and vice versa for rod bipolar cells ^49^. Therefore, we monitored the effect of the phosphomimetic T14E mutation on syntaxin-3b conformations. As a control, we also tested the effect of the so-called LE mutations (L165A, E166A), which bypass the requirement of Munc13-1 and induces a constitutively open form of syntaxin-1 ^15–17^. Interestingly, the phosphomimetic T14E mutant induced a 3-fold increase of molecules in the open conformation confirmed by the FRET efficiency histogram (Fig. 2D–E). As expected, the LE mutation on syntaxin-3b promoted an almost complete opening of the Habc domain with approximately 6-fold increase of molecules in the open conformation (Fig. 2D–E). Furthermore, the T14E mutation on syntaxin-3b increased the binding affinity with SNAP-25 by about 2-fold (Fig. 2F). Taken together, the phosphomimetic T14E mutation of syntaxin-3b induced the opening of the Habc domain, exposing the SNARE motif to allow interactions with SNAP-25, permitting SNARE complex assembly.

We next asked how efficient syntaxin-3b and its mutations form the ternary SNARE complex, composed of SNAP-25 and synaptobrevin. In agreement with the inefficient binding of SNAP-25 to wild-type syntaxin-3b, only about 50% of the molecules formed ternary SNARE complex, i.e., a shift to low FRET efficiency indicating the decrease in the percent closed population (Fig. 3D–E). In the presence of the phosphomimetic T14E mutation, the SNARE complex formation increased by 2-fold compared to the wild-type syntaxin-3b (Fig. 3D–E). To further corroborate the opening of syntaxin-3b by the phosphomimetic T14E mutation, we performed a pulldown assay using syntaxin mutants as a bait for SNAP-25 interactions. Consistent with our smFRET measurements, the T14E mutation on syntaxin-3b was able to recover more than 80% of SNAP-25 binding compared to the SNAP-25 levels of wild-type syntaxin-1a (Fig. 3F–G). As expected, the LE mutation fully recovered the binary t-SNARE complex formation with SNAP-25, indicating a complete shift to open conformation (Fig. 3F– G).

Since the SNARE chaperone Munc18-1 regulates the conformational switch of syntaxins by inducing an inhibitory closed conformation, we investigated the effect of Munc18-1 on syntaxin-3b conformations. As expected, Munc18-1 had little effect on the FRET efficiency histogram of wild-type syntaxin-3b since the wild-type already possesses a self-inhibiting closed conformation by itself (Fig. 4D–E). However, Munc18-1 was not able to completely induce a closed conformation for the syntaxin-3b T14E mutant, where about 30% of the molecules were still observed in an open conformation (Fig. 4D–E). To validate if the phosphomimetic T14E mutation on syntaxin-3b is capable of forming SNARE complex in the presence of Munc18-1, we performed a pulldown assay using the heterodimeric complex of Munc18-1 and syntaxin mutants as bait for SNAP-25 interactions. As anticipated, Munc18-1 bound tightly to syntaxin-1a and WT syntaxin-3b (Fig. 4F–G). Therefore, the binary t-SNARE complex formation with SNAP-25 decreased in the presence of Munc18-1 for both syntaxin-1a and WT syntaxin-3b by about 70% and 50%, respectively (Fig. 3G vs 4G). On the other hand, the binary t-SNARE complex formation of syntaxin-3b T14E mutant increased by almost 3-fold compared to the WT in the presence of Munc18-1 (Fig. 4F–G). This is likely due to the 50% decrease in the Munc18-1 levels of the pulldown assay using syntaxin-3b T14E mutant compared to the WT. Furthermore, Munc18-1 did not stay bound to syntaxin-3b LE mutant in our pulldown assay, consequently permitting complete formation of the binary t-SNARE complex with SNAP-25, comparable to the absence of Munc18-1 (Fig. 3G vs 4G).

Taken together, our single-molecule experiments, along with ensemble biochemical assays, revealed that syntaxin-3b is predominantly in a self-inhibiting closed conformation in contrast to syntaxin-1a, which adopts two conformations, i.e., an open and closed conformation, in its native state. Moreover, we demonstrated that the T14E mutation on syntaxin-3b, mimicking the phosphorylation by CaMKII, is capable of forming a binary t-SNARE complex with SNAP-25 in the presence of Munc18-1. In summary, we propose a model where phosphorylation on syntaxin-3b induces an opening of the regulatory Habc domain locked in an inhibitory closed conformation by Munc18-1, acting as a conformational switch to trigger SNARE complex assembly.

## Materials and methods

### Proteins: plasmids, expression, purification, and labeling

The cytoplasmic domains of syntaxin-1a (residues 1–265) and syntaxin-3b (residues 1–264) were fused with a C-terminal Avi-tag sequence (GLNDIFEAQKIEWHE) for biotinylation and an N-terminal TEV cleavable 6x-histidine tag for purification. The syntaxin-1a construct was cloned into the pTEV5 vector ^50^ and the syntaxin-3b construct was synthesized and inserted into pJ414 vector (ATUM, Newark, CA). Labeling sites were introduced by generating cysteine mutations using QuickChange Kit (Agilent, Santa Clara, CA) on a cysteine-free template. For intra-molecular smFRET measurements, two cysteine mutations were introduced on residues 35 and 249 for syntaxin-1a, and residues 37 and 248 for sytnaxin-3b construct. The phosphomimetic T14E mutation and the LE mutation on syntaxin-3b were also generated using QuickChange Kit (Agilent, Santa Clara, CA). Full-length SNAP-25 and the cytoplasmic domain of synaptobrevin-2 (residues 1–96) were cloned into the pTEV5 vector with an N-terminal TEV cleavable 6x-histidine tag.

All proteins were expressed in *E. coli* BL21 (DE3*) cells. The cells were grown in Terrific Broth (TB) medium at 37℃ until OD_600_ reached 0.8 and induced using 1 mM isopropyl β-D-1-thiogalactopyranoside (IPTG) overnight at 25℃. For *in vivo* biotinylation on syntaxin molecules, syntaxin constructs were co-expressed with the BirA gene engineered pACYC184 plasmid (Avidity, Aurora, CO) and induced with 1 mM IPTG in the presence of 0.1 mM biotin overnight at 25℃. Cells were harvested and resuspended in PBS buffer (50 mM NaH_2_PO_4_ pH 8.0, 300 mM NaCl, 0.5 mM TCEP). The cells were lysed by sonication and the lysate were clarified by using a Sorvall RC5C Plus centrifuge with an SS-34 rotor (ThermoFisher Scientific, Waltham, MA) at 20,000 rpm for 30 min. The supernatant was bound to 5 mL of Nickel-NTA agarose resin (Thermo Scientific, Waltham, MA) for 1 hour at room temperature. The resin bound with proteins was extensively washed with 50 mL PBS and eluted with PBS containing 400mM imidazole. For all proteins, the N-terminal 6x-histidine tags were cleaved by adding 2 mg of Tobacco Etch Virus (TEV) protease during dialysis overnight in 20 mM Tris pH 8.0, 50 mM NaCl, and 0.5mM TCEP. Proteins were further purified using HiTrap Q (GE Healthcare, Piscataway, NJ) anion exchange. The affinity tag for Munc18 was cleaved by 2 mg of TEV in PBS with 400 mM imidazole overnight at 4℃ and additional purification was performed using superdex 200 10/300 GL (GE Healthcare, Piscataway, NJ) gel filtration column in TBS buffer (20 mM Tris pH 7.5, 150 mM NaCl, 0.5 mM TCEP). The purity of all proteins was confirmed by SDS-PAGE.

Syntaxin molecules containing intra-molecular FRET labeling sites, i.e., double cysteine mutations, were stochastically labeled with Alexa 555 C_2_ and Alexa 647 C_2_ maleimide (Thermo Fisher Scientific, Waltham, MA) fluorescent dyes at 10x molar excess of the protein concentration in TBS overnight on a rotating platform at 4℃. The labeled proteins were separated from the free dyes on a homemade Sephadex G50 resin (GE Healthcare, Piscataway, NJ) column in TBS buffer.

### Single-molecule FRET measurements

To achieve single-molecule resolution, fluorescently labeled protein molecules were surface-tethered within a well-separated diffraction limited spot through biotin-streptavidin linkage as described previously ^51^. To prevent non-specific binding, the surface of the microscope slide was coated by polyethylene glycol (PEG) containing 0.1% (w/v) biotinylated-PEG. 0.1 mg/ml streptavidin was incubated for 5 min and rinsed with TBS buffer to allow surface-tethering of fluorescently labeled syntaxin molecules via the C-terminal biotinylation. Data was collected using the imaging buffer (1% (w/v) glucose, 20 mM Tris pH 7.5, 150 mM NaCl) in the presence of oxygen scavenger (20 units/mL glucose oxidase, 1000 units/mL catalase) and triplet-state quencher (100 μM cyclooctatetraene) to prevent rapid photobleaching and blinking of fluorophores.

Details on instrumental setup have been described elsewhere ^21, 52^. Briefly, a custom-built prism-based total internal reflection fluorescence (TIRF) microscope was used to conduct smFRET measurements at 10 Hz frame rate for about 200 sec until most of the dyes photobleached. The movies were collected using smCamera software kindly provided by Professor Taekjip Ha (Johns Hopkins University, Baltimore, MD). The data was processed using homemade scripts written in MATLAB (Mathworks, Natick, MA) and Igor Pro (WaveMetrics, Portland, OR). The FRET efficiency was calculated by E=I_A_/(I_A_+I_D_), where I_A_ and I_D_ are leakage corrected fluorescent intensities of acceptor and donor emissions, respectively ^53^.

### Fluorescence anisotropy binding assays

For the fluorescence anisotropy binding experiments described in Figures 1F and 2F, syntaxin molecules were labeled at the N-terminal primary amino group (-NH_2_) of the proteins using fluorescent dye Alexa 488 *N*-hydroxysuccinimide (NHS) (ThermoFisher Scientific, Waltham, MA) in PBS at pH 6.5 overnight at 4°C. The reaction was quenched and free biotins were separated using a homemade Sephadex G50 resin (GE Healthcare, Piscataway, NJ) column in TBS buffer (20 mM Tris pH 7.5, 150 mM NaCl, 0.5 mM TCEP). Anisotropy was measured with a Synergy H4 Hybrid Microplate Reader (BioTek, Winooski, VT) with xenon flash system at an excitation wavelength of 460 ± 40 nm and the emission wavelength of 528 ± 28 nm at room temperature. Alexa 488 labeled samples were diluted to a final concentration of about 0.5 μM in TBS for optimal signal to noise ratio. The labeled samples were mixed with unlabeled SNAP-25A at different concentrations and incubated at room temperature for about 15 min in a 96 well plate prior to data collection.

### Biotin-Neutravidin affinity pulldown assay

The pulldown assay was performed using neutravidin coated agarose resin (ThermoFisher Scientific, Waltham, MA). 100 μL bead volume (BV) of beads were equilibrated in TBS buffer (20 mM Tris pH 7.5, 150 mM NaCl, 0.5 mM TCEP) before adding about 500 μg of bait proteins labeled with biotin. Biotin molecules were labeled at the N-terminal primary amino group (-NH_2_) of the bait proteins using *N*-hydroxysuccinimide (NHS)-biotin (ThermoFisher Scientific, Waltham, MA) in PBS at pH 6.5. Biotinylation was quenched and free biotins were separated using a homemade Sephadex G50 resin (GE Healthcare, Piscataway, NJ) column in TBS buffer. Neutravidin beads coated with biotinylated proteins were transferred to 0.5 cm diameter disposable columns. The beads were washed with 10x BV of TBS to remove biotin-free proteins. 500 μg of SNAP-25A was incubated with syntaxin coated beads for 1 hour at room temperature to test the efficiency of binary t-SNARE complex formation. For pulldown assays in the presence of Munc18-1, syntaxin coated beads were incubated with about 300 μg of Munc18-1 for 1 hour following a 10x TBS washing step prior to SNAP-25A incubation (Fig. 4F). The beads were washed with 10x BV of TBS to ensure specific protein-protein interactions. The bead samples were transferred to a microcentrifuge tube and boiled at 95°C for 15 min with 2x SDS loading buffer to promote protein denaturation. Proteins were separated via SDS-PAGE electrophoresis and visualized by homemade Coomassie Blue staining. The gels were destained and scanned for analysis using ImageJ software (NIH, Bethesda, MD).

## Author contributions

C.G., J.F., and U.B.C., conceived this project. C.G., J.F., and U.B.C., designed, conducted, and analyzed experiments. C.G., J.F., U.B.C., drafted and revised the article. B.M., and G.H., interpreted and revised the article.

---

**Supplemental Figure 1.**
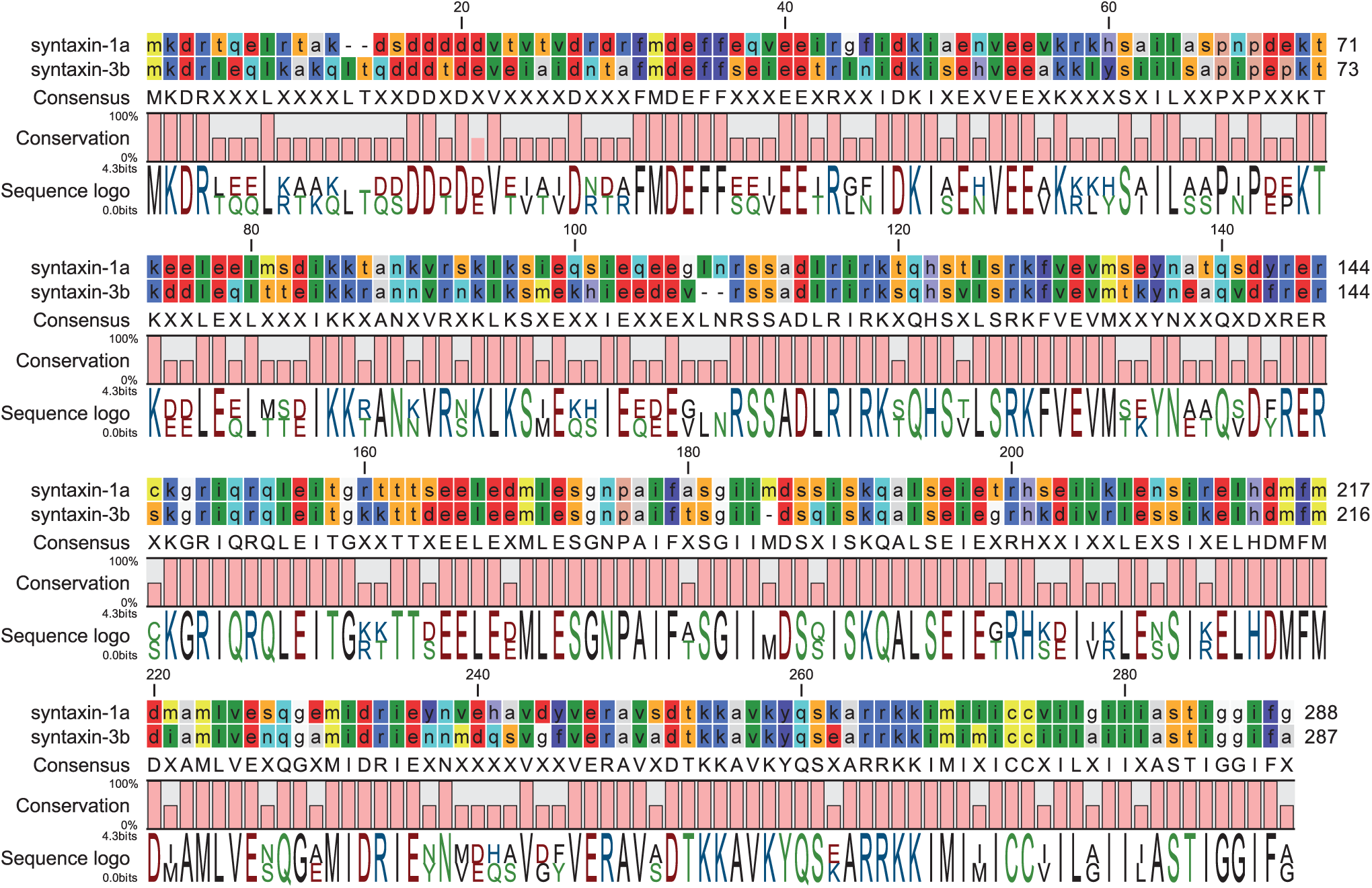
Sequence alignment of syntaxin-1a and syntaxin-3b. The amino acid colors are based on the RasMol color scheme according to the properties of the amino acid, i.e., polar residues are bright colors and non-polar residues are dark colors.

## References

1. Lagnado, L. & Schmitz, F. Ribbon Synapses and Visual Processing in the Retina. Annual Review of Vision Science vol. 1 (2015).

2. Heidelberger, R., Thoreson, W. B. & Witkovsky, P. Synaptic transmission at retinal ribbon synapses. Progress in Retinal and Eye Research vol. 24 686–720 (2005).

3. Moser, T., Grabner, C. P. & Schmitz, F. Sensory processing at ribbon synapses in the retina and the cochlea. Physiol. Rev. 100, 103–144 (2020).

4. Brunger, A. T., Leitz, J., Zhou, Q., Choi, U. B. & Lai, Y. Ca 2+ -Triggered Synaptic Vesicle Fusion Initiated by Release of Inhibition. Trends in Cell Biology vol. 28 631–645 (2018).

5. Südhof, T. C. & Rothman, J. E. Membrane fusion: Grappling with SNARE and SM proteins. Science vol. 323 474–477 (2009).

6. Brunger, A. T. et al. The pre-synaptic fusion machinery. Current Opinion in Structural Biology vol. 54 179–188 (2019).

7. Rizo, J. & Rosenmund, C. Synaptic vesicle fusion. Nat. Struct. Mol. Biol. 15, 665–674 (2008).

8. Sutton, R. B., Fasshauer, D., Jahn, R. & Brunger, A. T. Crystal structure of a SNARE complex involved in synaptic exocytosis at 2.4 Å resolution. Nature vol. 395 347–353 (1998).

9. Poirier, M. A. et al. The synaptic SNARE complex is a parallel four-stranded helical bundle. Nat. Struct. Biol. 5, 765–769 (1998).

10. Curtis, L. et al. Syntaxin 3B is essential for the exocytosis of synaptic vesicles in ribbon synapses of the retina. Neuroscience 166, 832–841 (2010).

11. Hays, C. L. et al. Simultaneous Release of Multiple Vesicles from Rods Involves Synaptic Ribbons and Syntaxin 3B. Biophys. J. 118, 967–979 (2020).

12. Liu, X., Heidelberger, R. & Janz, R. Phosphorylation of syntaxin 3B by CaMKII regulates the formation of t-SNARE complexes. Mol. Cell. Neurosci. 60, 53–62 (2014).

13. Curtis, L. B. et al. Syntaxin 3b is a t-SNARE specific for ribbon synapses of the retina. J. Comp. Neurol. 510, 550–559 (2008).

14. Misura, K. M. S., Scheller, R. H. & Weis, W. I. Three-dimensional structure of the neuronal-Sec1-syntaxin 1a complex. Nature vol. 404 355–362 (2000).

15. Wang, S. et al. Conformational change of syntaxin linker region induced by Munc13s initiates SNARE complex formation in synaptic exocytosis. EMBO J. 36, 816–829 (2017).

16. Gerber, S. H. et al. Conformational switch of syntaxin-1 controls synaptic vesicle fusion. Science (80-. ). 321, 1507–1510 (2008).

17. Dulubova, I. et al. A conformational switch in syntaxin during exocytosis: Role of munc18. EMBO J. 18, 4372–4382 (1999).

18. Wang, X. et al. Munc13 activates the Munc18-1/syntaxin-1 complex and enables Munc18-1 to prime SNARE assembly. EMBO J. 39, e103631 (2020).

19. Ma, C., Li, W., Xu, Y. & Rizo, J. Munc13 mediates the transition from the closed syntaxin-Munc18 complex to the SNARE complex. Nat. Struct. Mol. Biol. 18, 542–549 (2011).

20. Ma, C., Su, L., Seven, A. B., Xu, Y. & Rizo, J. Reconstitution of the Vital Functions of Munc18 and Munc13 in Neurotransmitter Release Downloaded from. Science *(80-. ).* **339**, 421–425 (2013).

21. Lai, Y. et al. Molecular Mechanisms of Synaptic Vesicle Priming by Munc13 and Munc18. Neuron 95, 591–607 (2017).

22. Rizo, J. & Xu, J. The Synaptic Vesicle Release Machinery. Annu. Rev. Biophys. 44, 339– 367 (2015).

23. Burkhardt, P., Hattendorf, D. A., Weis, W. I. & Fasshauer, D. Munc18a controls SNARE assembly through its interaction with the syntaxin N-peptide. EMBO J. 27, 923–933 (2008).

24. Colbert, K. N. et al. Syntaxin1a variants lacking an N-peptide or bearing the LE mutation bind to Munc18a in a closed conformation. Proc. Natl. Acad. Sci. U. S. A. 110, 12637– 12642 (2013).

25. Romero, P. et al. Sequence complexity of disordered protein. Proteins Struct. Funct. Genet. 42, 38–48 (2001).

26. Rickman, C., Meunier, F. A., Binz, T. & Davletov, B. High Affinity Interaction of Syntaxin and SNAP-25 on the Plasma Membrane Is Abolished by Botulinum Toxin E. J. Biol. Chem. 279, 644–651 (2004).

27. Morgans, C. W., Brandstätter, J. H., Kellerman, J., Betz, H. & Wässle, H. A SNARE complex containing syntaxin 3 is present in ribbon synapses of the retina. J. Neurosci. 16, 6713–6721 (1996).

28. Sherry, D. M., Mitchell, R., Standifer, K. M. & du Plessis, B. Distribution of plasma membrane-associated syntaxins 1 through 4 indicates distinct trafficking functions in the synaptic layers of the mouse retina. BMC Neurosci. 7, 1471–2202 (2006).

29. Foletti, D. L., Lin, R., Finley, M. A. F. & Scheller, R. H. Phosphorylated syntaxin 1 is localized to discrete domains along a subset of axons. J. Neurosci. 20, 4535–4544 (2000).

30. Shi, V. et al. Phosphorylation of Syntaxin-1a by casein kinase 2α regulates pre-synaptic vesicle exocytosis from the reserve pool. J. Neurochem. 156, 614–623 (2021).

31. Rickman, C. & Duncan, R. R. Munc18/syntaxin interaction kinetics control secretory vesicle dynamics. J. Biol. Chem. 285, 3965–3972 (2010).

32. Stein, A., Weber, G., Wahl, M. C. & Jahn, R. Helical extension of the neuronal SNARE complex into the membrane. Nature 460, 525–528 (2009).

33. Verhage, M. et al. Synaptic assembly of the brain in the absence of neurotransmitter secretion. Science (80-. ). 287, 864–869 (2000).

34. Wang, S. et al. Munc18 and Munc13 serve as a functional template to orchestrate neuronal SNARE complex assembly. Nat. Commun. 10, 69 (2019).

35. Stepien, K. P., Prinslow, E. A. & Rizo, J. Munc18-1 is crucial to overcome the inhibition of synaptic vesicle fusion by αSNAP. Nat. Commun. 10, 4326 (2019).

36. Rizo, J. & Südhof, T. C. The membrane fusion enigma: SNAREs, Sec1/Munc18 proteins, and their accomplices guilty as charged? Annu. Rev. Cell Dev. Biol. 28, (2012).

37. Brunger, A. T., Choi, U. B., Lai, Y., Leitz, J. & Zhou, Q. Molecular Mechanisms of Fast Neurotransmitter Release. Annu. Rev. Biophys. 47, 469–497 (2018).

38. Fernandez, I. et al. Three-dimensional structure of an evolutionarily conserved N-terminal domain of syntaxin 1A. Cell 94, 841–849 (1998).

39. Eisemann, T. J. et al. The sec1/munc18 protein vps45 holds the qa-snare tlg2 in an open conformation. Elife 9, e60724 (2020).

40. Lee, S. et al. Munc18-1 induces conformational changes of syntaxin-1 in multiple intermediates for SNARE assembly. Sci. Rep. 10, 11623 (2020).

41. Margittai, M. et al. Single-molecule fluorescence resonance energy transfer reveals a dynamic equilibrium between closed and open conformations of syntaxin 1. Proc. Natl. Acad. Sci. U. S. A. 100, 15516–15521 (2003).

42. Augustin, I., Rosenmund, C., Südhof, T. C. & Brose, N. Munc13-1 is essential for fusion competence of glutamatergic synaptic vesicles. Nature 400, 457–461 (1999).

43. Richmond, J. E., Davis, W. S. & Jorgensen, E. M. Unc-13 is required for synaptic vesicle fusion in C. elegans. Nat. Neurosci. 2, 959–964 (1999).

44. Aravamudan, B., Fergestad, T., Davis, W. S., Rodesch, C. K. & Broadie, K. Drosophila UNC-13 is essential for synaptic transmission. Nat. Neurosci. 2, 965–971 (1999).

45. Varoqueaux, F. et al. Total arrest of spontaneous and evoked synaptic transmission but normal synaptogenesis in the absence of Munc13-mediated vesicle priming. Proc. Natl. Acad. Sci. U. S. A. 99, 9037–9042 (2002).

46. Imig, C. et al. The Morphological and Molecular Nature of Synaptic Vesicle Priming at Presynaptic Active Zones. Neuron 84, 416–431 (2014).

47. Cooper, B. et al. Munc13-independent vesicle priming at mouse photoreceptor ribbon synapses. J. Neurosci. 32, 8040–8052 (2012).

48. Schmitz, F., Augustin, I. & Brose, N. The synaptic vesicle priming protein Munc13-1 is absent from tonically active ribbon synapses of the rat retina. Brain Res. 895, 258–263 (2001).

49. Campbell, J. R. et al. Phosphorylation of the Retinal Ribbon Synapse Specific t-SNARE Protein Syntaxin3B Is Regulated by Light via a Ca2 +-Dependent Pathway. Front. Cell. Neurosci. 14, 587072 (2020).

50. Rocco, C. J., Dennison, K. L., Klenchin, V. A., Rayment, I. & Escalante-Semerena, J. C. Construction and use of new cloning vectors for the rapid isolation of recombinant proteins from Escherichia coli. Plasmid 59, 231–237 (2008).

51. Choi, U. B., Weninger, K. R. & Bowen, M. E. Immobilization of proteins for single-molecule fluorescence resonance energy transfer measurements of conformation and dynamics. Methods Mol. Biol. 896, 3–20 (2012).

52. Choi, U. B. et al. NSF-mediated disassembly of on and off-pathway snare complexes and inhibition by complexin. Elife 7, 1–26 (2018).

53. McCann, J. J., Choi, U. B., Zheng, L., Weninger, K. & Bowen, M. E. Optimizing methods to recover absolute FRET efficiency from immobilized single molecules. Biophys. J. 99, 961–970 (2010).

